# Aberrant prefrontal beta oscillations predict episodic memory encoding deficits in schizophrenia

**DOI:** 10.1101/061291

**Authors:** Federica Meconi, Sarah Anderl-Straub, Heidelore Raum, Michael Landgrebe, Berthold Langguth, Karl-Heinz T. Bäuml, Simon Hanslmayr

**Affiliations:** Department of Developmental and Social Psychology, University of Padua, Italy; Department of Neurology, University of Ulm, Ulm, Germany; Department of Experimental Psychology, University of Regensburg, Regensburg, Germany; Department of Psychiatry and Psychotherapy, University of Regensburg, Regensburg, Germany; School of Psychology, University of Birmingham, Birmingham, UK

## Abstract

Verbal episodic memory is one of the core cognitive functions affected in patients suffering from schizophrenia (SZ). Although this verbal memory impairment in SZ is a well-known finding, our understanding about its underlying neurophysiological mechanisms is rather scarce. Here we address this issue by recording brain oscillations during a memory task in a sample of healthy controls and patients suffering from SZ. Brain oscillations represent spectral fingerprints of specific neurocognitive operations and are therefore a promising tool to identify neurocognitive mechanisms that are affected by SZ. Healthy controls showed a prominent suppression of left prefrontal beta oscillatory activity during successful memory formation, which replicates several previous oscillatory memory studies. In contrast, patients failed to exhibit such left prefrontal beta power suppression. Utilizing a new topographical pattern similarity approach, we further demonstrate that the degree of similarity between a patient's beta power decrease to that of the controls reliably predicted memory performance. This relationship between beta power decreases and memory was such that the patients' memory performance improved as they showed a more similar topographical beta desynchronization pattern compared to that of healthy controls. These findings suggest that left prefrontal beta power suppression (or lack thereof) during memory encoding is a possible biomarker for the observed encoding impairments in SZ in verbal memory. This lack of left prefrontal beta power decreases might indicate a specific semantic processing deficit of verbal material in patients with schizophrenia.

## Introduction

Any complex cognitive task requires a fine-grained balance between synchronized and desynchronized brain networks, which is regulated by oscillations (Buzsáki and Draguhn, 2004; DaSilva, 2013). An increasing number of studies show that this balance between synchronized and desynchronized networks is affected in patients with schizophrenia (SZ), who show markedly aberrant patterns of oscillatory activity in the lower (i.e. delta, theta, alpha, beta) and higher (gamma) frequency ranges (Uhlhaas and Singer, 2015). Arguably, oscillations reflect basic neurocognitive operations with different frequency bands representing different spectral “fingerprints” of these operations (Siegel, Donner, and Engel, 2012). A promising approach for SZ therefore is to identify the affected neurocognitive operations by isolating spectral fingerprints in specific tasks in patients, ultimately identifying targets for brain stimulation or pharmaceutical interventions. This approach has proven to be quite successful in the domains of perception (i.e. feature binding, Grützner, Wibral, Sun, Rivolta, Singer, Maurer, and Uhlhaas, 2013) and working memory (Haenschel, Bittner, Waltz, Haertling, Wibral, Singer, Linden, and Rodriguez, 2009), where deficits mainly in the beta and gamma frequency bands have been found (Uhlhaas and Singer, 2015). However, no study has so far investigated the oscillatory fingerprints of episodic memory formation deficits in SZ. This is quite remarkable given that episodic memory is particularly strongly affected in SZ (Schaefer, Giangrande, Weinberger, and Dickinson, 2013). We here aim to fill this gap and test whether specific oscillatory markers can explain cognitive deficits in verbal episodic memory encoding in SZ.

In the healthy brain, encoding of episodic memories is typically reflected by a well-orchestrated interplay of desynchronized low frequency activity in the neocortex (< 20 Hz) and synchronized theta/gamma activity in the hippocampus (Hanslmayr, Staresina, and Bowman, 2016). Within this framework, the active engagement of specific cortical modules is mediated by a pronounced desynchronization in the low frequencies which allow these modules to represent information in a rich manner (Hanslmayr, Staudigl, Fellner, 2012). For instance, formation of non-verbal memories has been associated with cortical theta and gamma power increases as well as fronto-temporal phase-amplitude coupling (Mölle, Marshall, Fehm, and Born, 2002; Lega, Jacobs and Kahana, 2012; Hanslmayr, Spitzer and Bäuml, 2009; Osipova, Takashima, Oostenveld, Fernández, Maris, and Jensen, 2006; Friese, Köster, Hassler, Martens, Trujillo-Barreto, and Gruber, 2013; Köster, Friese, Schone, Trujillo-Barreto, and Gruber, 2014; Lega, Burke, Jacobs, and Kahana, 2016). Concerning beta oscillations a different picture arises. Cumulating evidence revealed that when participants are required to intentionally encode a list of words (Hanslmayr and Staudigl, 2014; Hanslmayr, Volberg, Wimber, Raabe, Greenlee, and Bäuml, 2011; Hanslmayr et al., 2012) or to incidentally encode a list of words via accessing their semantic content (Hanslmayr et al., 2009, 2011; Fellner, Bäuml and Hanslmayr, 2013) successful encoding of verbal material into episodic memory is reflected by a pronounced beta power decrease (13-20 Hz) in the left inferior prefrontal cortex (IFG), as revealed by MEG (Meeuwissen, Takashima, Fernández, Jensen, 2011), simultaneous EEG-fMRI (Hanslmayr et al., 2011) and rTMS-EEG (Hanslmayr, Matuschek, Fellner, 2014) studies. Left prefrontal beta power decreases have also been shown to reflect the level of processing of semantic material, thereby stronger power decreases indicate deeper, i.e. semantic processing and therefore better memory performance (Hanslmayr et al., 2009). Taken together these results suggest that the oscillatory signature of memory formation varies with the neural processes involved during memory encoding, and that the beta power decreases in the left prefrontal cortex reflect a deeper (i.e. more semantic) level of processing, especially for verbal material. Interestingly, a recent TMS study suggested a link between aberrant prefrontal beta oscillations and verbal memory deficits in SZ patients, however, since brain oscillations in this study were not measured during a memory task little information could be gleaned as to which memory process is affected in SZ (Ferrarelli, Sarasso, Guller, Riedner, Peterson, Bellesi, … and Tononi, 2012). Functional magnetic resonance imaging studies, on the other hand, demonstrated that SZ patients show less activation in the left IFG compared to controls during memory formation (Achim and Lapage, 2005). These findings therefore render prefrontal beta oscillations a promising marker for specific processing deficits in episodic memory encoding in SZ. A further advantage of prefrontal beta oscillations is that they can be robustly measured with non-invasive EEG, as opposed to the hippocampal theta-gamma activity, which does not lend itself easily to scalp EEG recordings.

In the present study we test the hypothesis that prefrontal beta oscillations indicate deficits in memory encoding in SZ patients. To this end we investigate oscillatory correlates of memory encoding in the so-called subsequent memory paradigm (Paller and Wagner, 2002). Therein, oscillatory activity was recorded with EEG during encoding of short lists of words. Items were back-sorted according to whether they were successfully retrieved or not in a delayed free recall test. Oscillatory responses that distinguish between later retrieval and later forgetting are referred to as subsequent memory effects (SME; see Figure 1A). These subsequent memory effects were contrasted between patients diagnosed with SZ and healthy controls.

**Figure 1.**
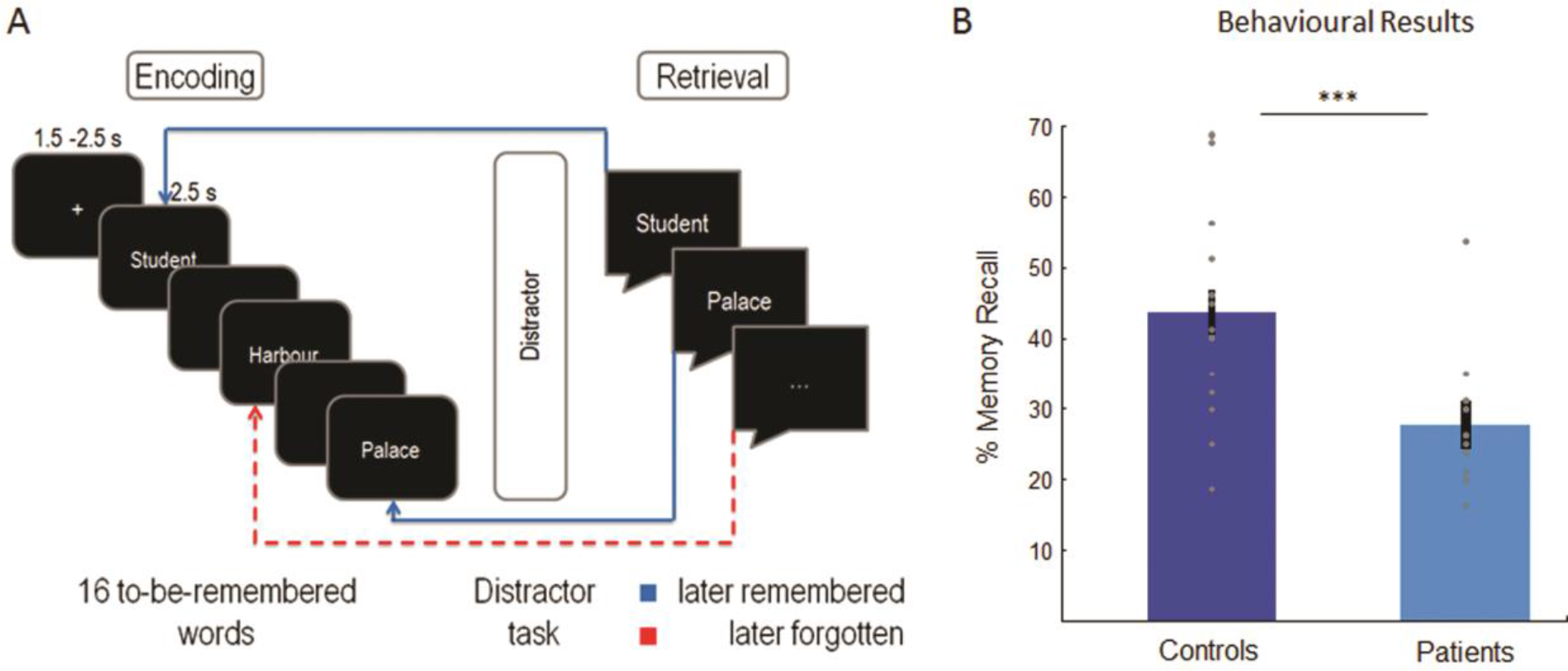
Experimental procedure and behavioral results. (A) The experimental procedure and logic of the subsequent memory effect (SME) analysis is shown. Participants encoded lists of 16 words and freely recalled as many words as possible in the retrieval phase, after a distractor task. (B) Average memory performance and individual data points of both groups of participants are shown (note the outlier in the SZ group). Error bars represent standard errors.

## Methods

### Patients

Twenty-four patients were initially recruited from the local psychiatric clinic (Regensburg, Germany). The data of four patients were excluded because too few trials remained after artifact rejection and after splitting trials into subsequently remembered (hits) and subsequently forgotten (misses), thus not allowing for a meaningful subsequent memory contrast (i.e. <13 trials; see Hanslmayr et al., 2009 for a justification for this trial number). The data of the first three patients were excluded because the paradigm was changed afterwards (i.e. lists were shortened because memory performance was too low); the data of two further patients were excluded because of incomplete datasets. The remaining sample consisted of 15 patients, all of whom suffered from schizophrenia of either the paranoid-hallucinatory (n = 10; ICD-10, F20.0) or hebephrenic subtype (n = 5; ICD-10, F20.1). The patients had a mean duration of illness of 66.6 months (SD, 71.02). 14 patients received atypical neuroleptics only; one patient received no medication at all. Mean chlorpromazine equivalent dosage (CPZ, Woods, 2003) was 559.63 mg/d (SD, 557.29). Two patients received antidepressant medication (selective serotonin reuptake inhibitors, or tetracyclic antidepressants). No patient had any neurological diseases, additional psychiatric diseases, or medication with tranquillizers. Current clinical symptoms were assessed with the positive and negative syndrome scale in all but one patient for whom this data is missing (PANSS, Kay, Flszbein and Opfer, 1987; N=14; mean score, 73.6; SD, 14.3).

### Controls

Twenty healthy volunteers served as control subjects, matched according to gender (4 females), educational level (M = 3 where 0: no graduation–4: university entrance diploma), handedness (all right-handed), amount of cigarettes smoked per day (M = 9.24, SD = 7.62), and premorbid IQ (M = 103.53, SD = 8.62) measured with the “Mehrfachwahl-Wortschatz-InteUigenztest” Lehrl, 2005). Although healthy volunteers and patients were both groups of young adults, healthy volunteers were on average slightly younger than patients (M = 24.35, SD = 4.55; t(30) = −2.53 p = 0.02). None of the participants had any history of neurological or psychiatric diseases or medication of any kind. Three healthy volunteers were discarded from analysis because of incomplete data, leaving a sample of 17 full datasets of control participants.

Demographic and IQ data and the number of cigarettes smoked per day for the final sample of controls and patients are plotted in Table 1. All of the participants had normal or corrected-to normal vision and were German native speakers. All gave written informed consent. The protocol was approved by the local ethical review board.

**Table 1.** Demographic data of control participants and patients. The table reports mean values and standard deviations (in brackets) of age, cigarettes per day and verbal IQ. The number of male and female participants and their handedness are also reported. The values of the statistical tests in the last row of the table show that the two groups are matched for cigarettes smoked per day, verbal IQ, gender and handedness. Median values are reported for educational values.

|  | Gender (m/f) | Handedness<br>RH/LH | Education | Age | Cigarettes<br>per day | Verbal IQ |
| --- | --- | --- | --- | --- | --- | --- |
| Controls (N = 17) | 13/4 | 17/0 | 3 | 24.35 (4.55) | 9.24 (7.62) | 103.53 (8.62) |
| Patients (N = 15) | 10/5 | 14/1 | 2.5 | 29.2 (6.26) | 13.33 (12.28) | 100.13 (17.45) |
| Stat. Difference | $\chi^2 = .379$; $p = .7$ | $\chi^2 = 1.17$; $p = .45$ | $Z = 1.05$; $p = .29$ | $t(30) = -2.53$ $p = .02$ | $t(22.82) = -1.062$ $p = .3$ | $t(30) = .711$ $p = .48$ |

### Stimuli and Procedure

Stimuli were comprised of 80 words drawn from the MRC Psycholinguistic Database (Coltheart, 1981), translated into German. The words were split into 5 lists of 16 words each. The lists were matched according to word frequency (*M*: 52.95, *SD:* 51.12), number of letters (*M*: 5.36, *SD:* 1.15), syllables (*M*: 1.69, *SD:* 0.54), concreteness (mean, 542.9; SD, 42.5), and imagability (*M*: 563.24, *SD:* 32.3). Stimuli were presented in white font on a black background of a 17’’ computer screen with a refreshing rate of 70 Hz. Participants performed a verbal long-term memory task, which was comprised of three phases: (i) encoding; (ii) distracter task; and (iii) free recall. The task was programmed using Presentation (© Neurobehavioral Systems). In the encoding phase, 16 words were sequentially presented in white font on a black background. Each word was presented for 2.5 seconds, preceded by a fixation cross with a variable duration of 1.5-2.5 seconds. Participants were instructed to try to keep the presented words in memory as they will be required to recall them later. Half-way through the list a reminder was presented prompting the participant to try to keep the presented words in memory. After the encoding phase a distracter task was carried out for 30 seconds, requiring the participants to rate the attractiveness of faces. The purpose of the distracter task was to minimize the contribution of working memory. Thereafter, an instruction was presented asking the participant to try to recall as many words as possible. Participants were given 1 minute to recall the 16 words. Then the next study-test cycle started with a new list of words. Overall, 5 such study-test runs were conducted. The structure of one study-test run is depicted in Figure 1. Most subjects also participated in a separate attention paradigm which is reported elsewhere (Hanslmayr, Backes, Straub, Popov, Langguth, Hajak, Bäuml, and Landgrebe, 2013).

### EEG recording and analysis

The EEG was recorded from 62 channels, distributed equidistantly over the scalp and placed on an elastic cap (EasyCap) referenced to Cz. The EEG was re-referenced offline to the average reference. One additional external electrode was placed below the left eye to record vertical EOG. Impedance of the electrodes was kept below 20 KΩ. EEG and vertical EOG signals were amplified between 0.1-100 Hz with a notch filter at 50 Hz and digitized at a sampling rate of 500 Hz.

*EEG*—*analysis*. EEG data was analyzed with MATLAB (©The Mathworks, Munich, Germany) using the open-source FieldTrip toolbox (Oostenveld, Fries, Maris, Schoffelen, 2011; http://fieldtrip.fcdonders.nl/) and in-house Matlab routines. EEG data were first segmented into epochs of 6 s, starting 2 seconds before word onset. The epoched data was visually inspected to discard large artifacts from further analysis. Remaining preprocessing steps included Independent Component Analysis for ocular artifacts correction and re-referencing to average reference. After removing trials which were contaminated by eye and muscle artefacts, an average of 30.5 trials (range: 13-52) remained for later remembered condition and 41.5 trials (range: 22-59) for later forgotten condition for the control subjects. For patients, an average of 19.4 trials (range: 13-39) and 52.8 (range: 38-61) for later remembered and later forgotten conditions remained for data analysis, respectively.

The segmented epochs (6 s) were then subjected to a wavelet transformation using Morlet wavelets (with a width of seven cycles) as implemented in fieldtrip to extract time-frequency characteristics optimally for each frequency. The frequency range of interest was 1-40 Hz with a time resolution of 50 ms. Power values were calculated for each single trial, and averaged across trials within the two conditions (later remembered, later forgotten). These power values were then transformed to relative power changes from baseline using an interval of −0.75 to −0.25 s before word onset (according to the formula: 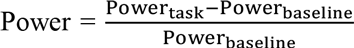) as implemented in fieldtrip). We finally subtracted power values for later forgotten words from power values for later remembered words to compute the subsequent memory effect (i.e., SME) for each participant.

We confined statistical analysis to the left hemisphere in the frequency range including alpha and beta bands (8-20 Hz), according to our hypothesis. To control for multiple comparisons nonparametric randomization tests were conducted (using 1000 permutations) averaging the power data across all channels of the left hemisphere, along the time-window of word presentation (2.5s). We performed this analysis first separately for each group (i.e. controls and patients) and then directly contrasted the SMEs between the two groups.

### Topographical similarity analysis

We computed a topographical similarity measure between patients and control participants in order to quantify the degree to which an EEG pattern in a given patient diverged in their SME from control subjects. In order to obtain individual patient topographies, power values were averaged over a 1 second long time-window centered at the point of maximal power decreases (later remembered-later forgotten) in each patient in a frequency window between 8 to 20 Hz (i.e., individual patients’ SMEs). Correlation indexes (i.e., Pearson’s *rs*) were calculated between these individual topographies and the topography of the grand average of controls. Power values of the topography of controls were averaged over a one second long time window centered at the point of maximal power decreases as shown by t-values obtained with statistical analysis depicted in Figure 2 (middle panel).

### Source Analysis

For source reconstruction, a frequency-domain adaptive spatial filtering dynamic imaging of coherent sources (DICS) algorithm (Groß, Kujala, Hamalainen, Timmermann, Schnitzler, Salmelin, 2001), as implemented in fieldtrip was applied. As source model a standardized boundary element model was used, which was derived from an averaged T1-weighted MRI dataset (MNI, www.mni.mcgill.ca). Source analysis was carried out for the frequency range which revealed significant results on the scalp level.

### Statistical analysis

Mean proportions of accurate recall words was averaged across blocks and lists and compared between patients and controls via an independent samples t-test. For correlations, Pearson’s correlations were calculated unless stated otherwise.

## Results

### Behavioural results

As expected, controls recalled significantly more words in the verbal memory task compared to patients diagnosed with SZ (see Figure 1B; *t(30)* = 3.85, *p* = .001). On average controls recalled 44%, (SD α13.7), whereas patients recalled only 28 % (SD ± 8.87), even though there is one patient who performed more than 2 SD above the average (53.75% see Figure 1); no outliers have been registered in healthy participants. No significant correlations between memory performance and CPZ (*r* = 0.21; *p* = 0.24), duration of illness (*r* = 0.20; *p* = 0.25) or age of the patients (*r* = −0.17; *p* = 0.18) were obtained either with or without exclusion of the outlier, suggesting that these variables were not the main contributors to the observed memory deficits in SZ.

### EEG results

In a first step, subsequent memory effects (SMEs) were analysed for healthy controls. A cluster analysis over the whole scalp for control participants revealed a significant SME in a cluster of left fronto-temporal electrodes (*p_corr_* = 0.023) in the beta frequency range (15-16 Hz) from 0.5 to 1.5 s after stimulus onset (Figure 2A). This beta oscillatory SME was due to a power decrease for subsequently remembered compared to subsequently forgotten words (Figure 2B), replicating previous findings (Hanslmayr et al., 2009, 2011, 2014; see also Kim, 2011 for a meta-analysis reporting left frontal SMEs for verbal information). Based on this result, further analysis were restricted to electrodes in the left hemisphere for both groups. Brain oscillatory SMEs are plotted in Figure 2 for both controls and patients. A significant SME was observed in control participants by contrasting oscillatory activity for subsequently remembered and subsequently forgotten trials in the time window during word presentation in a frequency range between 8 and 20 Hz and considering all electrodes of the left hemisphere (Figure 2A middle and right panels). Source level analysis revealed that this effect was mainly driven by sources in the left inferior frontal gyrus (MNI: -60 21 6; BA 45, see Figure 2B). Overall, these results show a normal subsequent memory effect in the control participants, i.e. stronger beta power decreases in the left prefrontal cortex during successful verbal memory formation confirming previous results (Meeuwissen et al., 2011; Hanlsmayr et al., 2009, 2011, 2014). In contrast, patients failed to show a significant SME (*p_corr_* >.7; Figure 2C). Moreover, a direct comparison in the left hemisphere of the beta SMEs between patients and controls in a 600 ms time-window (0.4-1s) between 10 and 18 Hz revealed a significant difference between patients and controls (*p_corr_*< 0.05) showing that the beta SME in the left hemisphere for controls was stronger compared to patients (Figure 2D).

**Figure 2:**
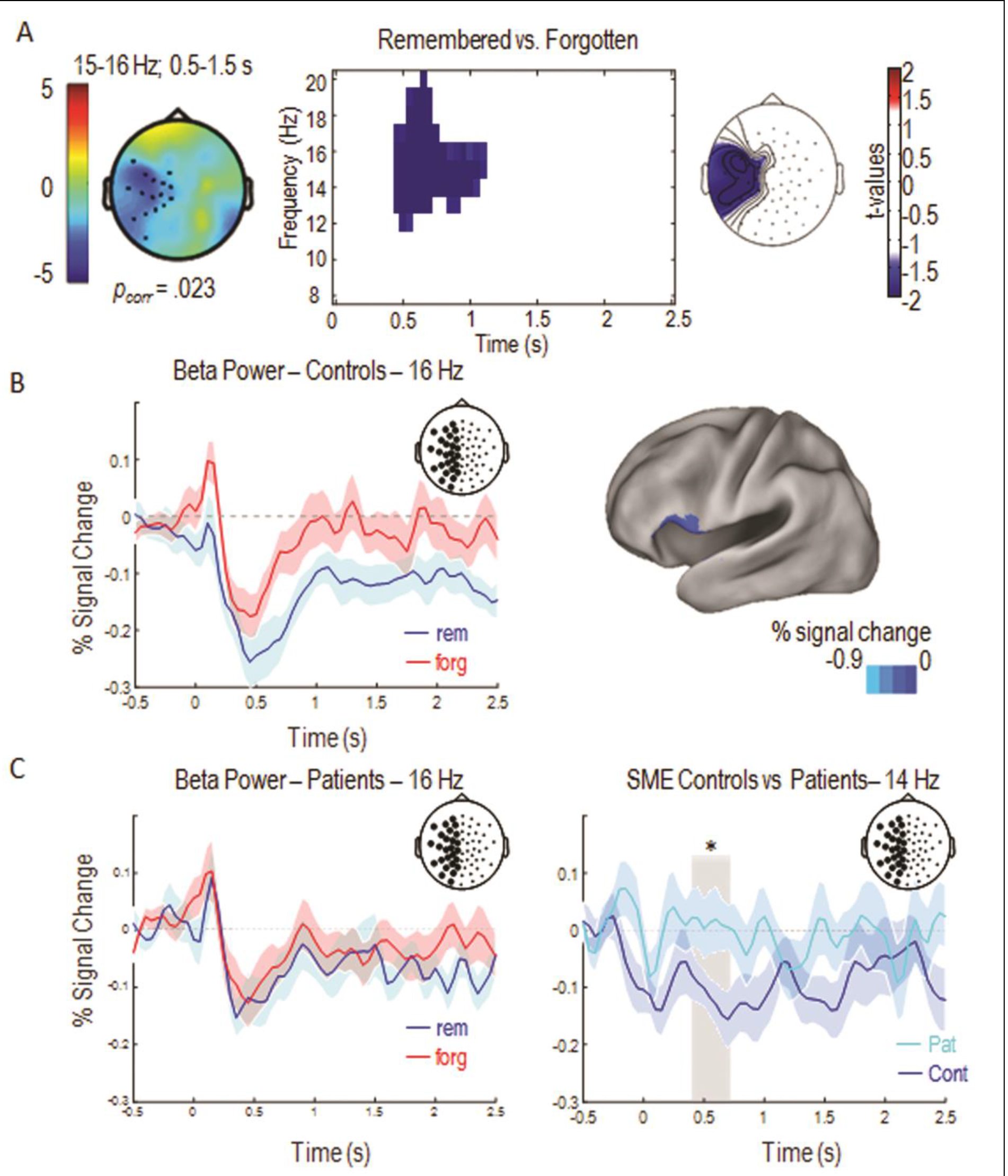
Subsequent Memory Effects. (A) The left panel shows the topography of the subsequent memory effect for controls (SME; 15-16 Hz, 0.5-1.5 s). Dots represent a significant cluster of electrodes, blue color indexes power decreases for later remembered words compared to later forgotten ones. The central panel shows a time-frequency plot of significant SMEs (t-values) averaged over all electrodes of the left hemisphere (central panel). The right panel shows the topography (t-values) of the beta SME. (B) The time-course of the power decrease at 16 Hz for subsequently remembered (blue) and forgotten (red) words of healthy controls is shown on the left. Source localization of the SME observed at the scalp level in healthy controls (15 Hz, 0.5 - 1.5 s; MNI: -60 21 6; right panel). (C) The time-course of the power decrease at 16 Hz for subsequently remembered (blue) and forgotten (red) words in SZ participants (left panel) and the time course of the contrast between controls’ (blue) and patients’ (light blue) SMEs (right panel); light grey rectangle represents the time-window (0.4-1 s) where the difference between SMEs has been observed. Shaded areas represent mean standard error. Circles in the pictures of the electrode positions on the scalp represent the averaged electrodes of the left hemisphere where SMEs and the contrast between groups’ SMEs have been observed.

### Topographical similarity results

Although the above results suggest that, on average, patients lack a left frontal beta SME compared to controls, inspection of individual datasets revealed that there was quite some variance with some patients actually showing left frontal beta power decreases quite similar to those obtained in controls (see Supplementary Figure 1). If these left frontal beta power decreases are functionally linked to the deficits in the patients’ memory performance, we should be able to predict the patients’ memory performance from their beta SME patterns. In order to formally test this hypothesis we computed a topographical similarity analysis between the SME patterns in patients and controls. To this end, we calculated correlation indexes (i.e., Pearson’s *rs*) between the individual patient SME topographies and the average SME topography obtained from the control subjects (Figure 3A; see Materials and Methods) and correlated this similarity index with memory performance. For this correlation analysis the outlier in memory performance (see Figure 1) was excluded (see Supplementary Figure 1 for the individual patient topographies). A remarkably strong positive correlation was observed between the patients’ topographical similarity values and memory performance (*r* = .683, *p* = .004). In other words, the more similar the beta SME pattern of a patient was to the pattern observed in the control group, the better the patient performed in the memory task. The correlation results are depicted in Figure 3. Critically, this correlation still holds when including the outlier and calculating Spearman's correlation (*ρ* = 0.635 *p* = 0.005), and when controlling for several critical aspects such as medication (*r* = .667 *p* = .006) duration of illness (*r* = .669 *p* = .006), the alpha/beta peak frequency where the strongest power decrease was observed (*r* = .530 *p* = .031), age of patients (*r* = .639 *p* = .009) and the individual number of trials (*r* = .684 *p* = .005), which is lower for patients than controls, due also to the lower accuracy rate. It is important to report that individual patients’ SMEs did not correlate with patients’ memory performance (*r* = -.06 *p* = .85), which suggests that the topographical distribution of beta power decreases (i.e., in the frontal regions of the left hemisphere), not the overall amount of power decreases, explains the variance in memory performance.

**Figure 3:**
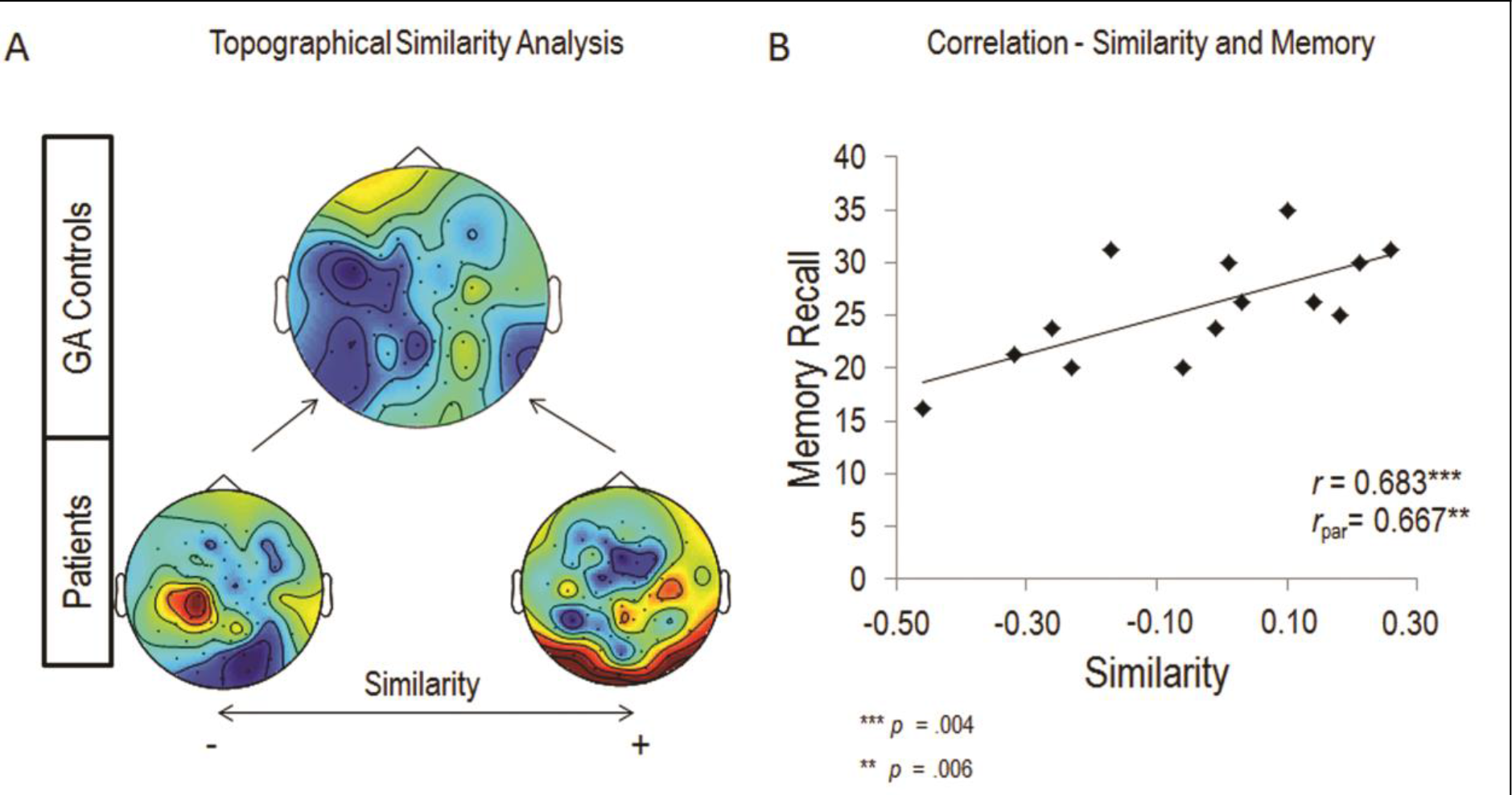
Topographical similarity analysis. (A) The grand average SME topography of the healthy participants (top), is shown together with the least (lower left) and the most similar topography (lower right) of the patients; blue represents power decreases, red represents power increases for later remembered words relative to later forgotten ones. (B) The scatterplot depicts the patients’ memory performance against the topographical similarity values (i.e. similarity between patients’ individual topographies and the grand average of healthy participants’ topographies; outlier excluded).

### Theta and gamma frequency bands

We also tested for SMEs in the gamma or theta frequency bands. In line with previous findings showing gamma synchronization associated with incidental encoding of non-verbal rather than intentional encoding of verbal material we did not observe any gamma activity (min *p* = .67). We did not either observe any theta synchronization after Montecarlo permutation (min *p* = .3). These null-results with regards to theta and gamma are largely consistent with our and other previous SME studies reporting that alpha/beta power decreases are the most robust signatures of memory encoding in EEG (Hanslmayr et al., 2012).

## Discussion

In the present study we show that deficits in episodic memory encoding of verbal material in patients suffering from SZ are associated with aberrant left prefrontal beta oscillations. This conclusion is based on two findings (i) SZ patients did not show the typical left prefrontal subsequent memory effect compared to controls; (ii) the degree to which an individual patient’s memory was impaired was strongly related to the dissimilarity between their beta oscillatory memory pattern and that of controls. This latter finding is especially important because it suggests that aberrant left prefrontal beta oscillations reflect memory encoding deficits in SZ.

On a behavioral level, healthy participants could recall significantly more words than patients, which is in good agreement with a recent meta-analysis showing that episodic memory is especially impaired in SZ (Schaefer et al., 2013). In the EEG, we found that successful encoding of verbal material in controls was indicated by a left inferior prefrontal beta power decrease, which replicates a number of previous studies (see Hanslmayr et al., 2012; Hanslmayr, and Staudigl, 2014, for reviews). However, patients with SZ failed to show such a typical SME pattern, which confirms our initial hypothesis. This result is in good agreement with previous fMRI studies showing decreased BOLD signal in the left inferior prefrontal cortex for SZ compared to controls during memory encoding (Achim and Lapage, 2005), as well as with a previous TMS study showing a link between aberrant prefrontal beta oscillations in SZ and memory (Ferrarelli et al., 2012). Importantly, our study goes beyond these previous findings in measuring beta oscillations during a memory encoding task and links these oscillations directly to memory impairments in SZ.

Left prefrontal beta power decreases have not only been linked to memory encoding, but also to semantic/conceptual processing of verbal material (Singh, 2012; Wang, Jensen, Van den Brink, Weder, Schoffelen, Magyari, … and Bastiaansen, 2012). Since processing words in a semantic/conceptual fashion has been shown to be a particularly efficient (i.e. deep) memory encoding strategy (Craik and Lockhart, 1972), left prefrontal beta power decreases likely indicate the depth of such semantic processing and therefore predict whether a given word is being later remembered or not (Hanslmayr and Staudigl, 2014). Indeed, instructing subjects to focus on semantic aspects of items during memory encoding leads to stronger left prefrontal beta power decreases, compared to less semantic encoding instructions (Hanslmayr, Spitzer, and Bäuml, 2009). Given this background, our findings suggest that patients might not be able to process items in such a deep, semantic fashion as controls, thereby limiting their encoding abilities. This explanation fits with the fact that conspicuous semantic processing (i.e. disorganized speech, neologisms, etc.) is a core diagnostic criteria of SZ. For instance, previous studies attempted to explain how schizophrenic patients have difficulties in verbal communication. Patients with SZ show reduced engagement of brain areas involved in the semantic organization of speech and grammar articulation (Kircher, Liddle, Brammer, Williams, Murray, and McGuire, 2001; Kircher, Oh, Brammer, and McGuire, 2005) and abnormal hemodynamic activity in response to either semantic associations (Kuperberg, Deckersbach, Holt, Goff, and West, 2007) and in semantic decision tasks (Sommer, Ramsey, and Kahn, 2001). Psycholinguistic studies of semantic processing in schizophrenia have further disclosed difficulties of patients with SZ in using context or combinatorial linguistic information, such as syntactic rules, to interpret appropriately the meaning of single words or of the whole sentences (see Kuperberg, 2010a,b for reviews). Impaired semantic processing in SZ is also indicated by decreased amplitudes of the N400 (Jackson, Foti, Kotov, Perlman, Mathalon, and Proudfit, 2014), which has been shown to strongly correlate with left prefrontal beta power decreases (Wang et al., 2012). Together, verbal memory encoding deficits, possibly due to impaired semantic processing, observed in SZ patients seem to be associated to weaker left prefrontal beta power suppression. Therefore, although the sample size of the SZ group was small, the here presented results suggests a new target for therapeutic interventions tailored to alleviate fundamental problems in memory encoding in SZ. A previous combined rTMS-EEG study in healthy subjects has shown that left prefrontal beta oscillations can be modified via rhythmic transcranial magnetic stimulation (rTMS) and that this modulation affects memory encoding (Hanslmayr et al., 2014). An interesting future application of rTMS therefore is to develop protocols in order to suppress left prefrontal beta power and to increase memory encoding as a result thereof. Alternatively, transcranial alternating current stimulation could be used to target left prefrontal beta oscillations in a suppressive manner. Although these protocols are yet to be developed and tested in healthy subjects, our study might inspire the development of such neuro-stimulation protocols, which are being increasingly recognized as treatment options for psychiatric conditions (Downar, Blumberger, Daskalakis, 2016). Moreover, our results might stimulate the development of pharmacological intervention that target neurotransmitters which are involved in the generation of beta oscillations, i.e. glutamate, NMDA receptors and GABAa receptors (Traub, Bibbig, LeBeau, Buhl, and Whittington, 2004; Yamawaki, Stanford, Hall, and Woodhall, 2008).

It is important to point out some potential caveats of our study, which were inevitably introduced by the clinical setting (i.e. patient availability, memory performance). For instance, the sample of patients was comprised of two different subtypes of SZ (i.e. 10 suffered from paranoid-hallucinatory and 5 from hebephrenic subtype). Furthermore the duration of illness and medication they were receiving at the time of data collection did vary considerably between patients; and patients were on average older than controls. However, arguably none of these aspects was the driving force in the observed correlation between beta similarity and memory performance as shown by the partial correlation analyses, further strengthening the argument that a lack of prefrontal beta suppression represents a core neurophysiological correlate of memory encoding impairments in schizophrenia. Finally, we cannot rule out a possible involvement of other oscillatory mechanisms which are not so readily accessible to non-invasive EEG (i.e. theta/gamma oscillations in the hippocampus), which is an important question for future studies to address.

## Conclusions

We investigated the oscillatory signature of episodic verbal memory formation in groups of healthy and schizophrenic participants. In line with previous studies, we observed a beta desynchronization SME in the left inferior frontal gyrus in healthy controls. Such a beta signature of episodic memory formation was absent in SZ patients. However, patients who showed a more similar topographical pattern of beta desynchronization compared to that of healthy controls, also performed better in the memory task. Together, these findings demonstrate that aberrant left prefrontal beta oscillations are closely linked to the memory impairments in SZ in a verbal memory task. These findings might ultimately inspire the development of neuro stimulation and pharmacological therapies which target left prefrontal beta oscillations in order to improve memory in SZ.

## Acknowledgements

The authors thank the patients for their willingness in participating in the experiments. This research was supported by grants from the Deutsche Forschungsgemeinschaft and the European Research Council awarded to SH (HA 5622/1-1 and 647954). The authors declare no competing financial interests.

## Supplementary Information

**Supplementary Figure 1.**
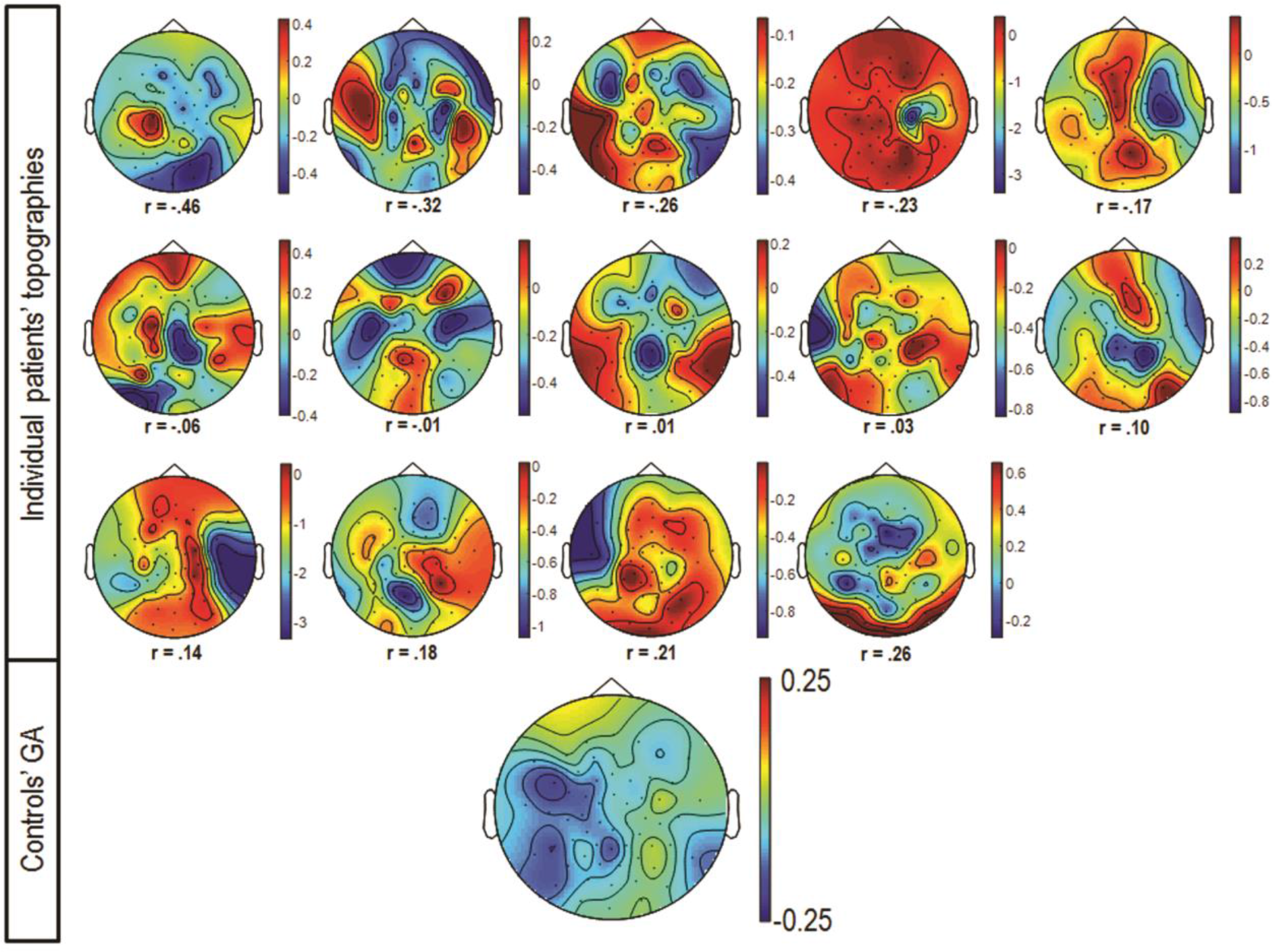
Individual patients’ topographic distribution of maximum power decrease associated with subsequent memory effect (SME) observed along 8-20 Hz frequency range in 1 s time-window (top panel; outlier not included; similarity values are presented by Pearson’s *rs*). Grand average of controls’ topographic distribution of maximum SME.

